# Investigations into fission yeast chromosome size determinants

**DOI:** 10.1101/2025.09.15.676266

**Authors:** Pei-Shang Wu, Todd Fallesen, Frank Uhlmann

## Abstract

Mitotic chromosome dimensions differ between organisms, and they differ within a species between developmental stages. The physiological determinants of chromosome size remain poorly understood. Here, we investigate chromosome size determinants in the fission yeast *Schizosaccharomyces pombe*. Super-resolution microscopy and semi-automated measurements reveal that cell or nuclear volume in interphase, or the time spent in mitosis (both previously proposed chromosome size determinants), have little influence on resultant chromosome dimensions. Instead, levels of the chromosomal condensin complex affect chromosome size, with increasing condensin levels resulting in more compact, shorter and thinner, chromosomes. These observations inform our understanding of how chromosome dimensions are controlled in an organism. They suggest that a chromosome-intrinsic mechanism sets chromosome size, more so than the environment in which chromosomes find themselves in.

## Introduction

Mitotic chromosome dimensions vary between eukaryotic species, which can harbour genomes of vastly different sizes (Flemming 1882; Sumner 2003; Kramer et al. 2021). Even in species with similar genome content, chromosome dimensions vary. E.g. the budding yeast *Saccharomyces cerevisiae* and the fission yeast *Schizosaccharomyces pombe* harbour approximately similarly sized genomes, divided amongst 16 short and thin or 3 long and thick chromosomes, respectively (Kakui et al. 2022). What defines the correct chromosome width in both species remains unknown. Another pair of organisms with similarly sized genomes are the closely related Chinese and Indian muntjacs. The DNA is distributed amongst 46 short and thin, or only 6 much longer and also much wider chromosomes in these two species (Wurster and Benirschke 1967; Wurster and Benirschke 1970). Again, how the Chinese and Indian muntjac chromosomes adopt their correct respective widths remains unknown.

Chromosome dimensions also vary within a species, e.g. accompanying the reduction of cell size by cleavage divisions during early embryonic development of multicellular organisms. The idea that chromosome size follows the size of the cell nucleus during early development of sea snails reaches back over 100 years ago (Conklin 1912). In the nematode *C. elegans*, chromosomes start relatively long at the one-cell stage, and they gradually shorten over the course of the first 8 cleavage divisions (Hara et al. 2013). The increased DNA density in progressively smaller interphase nuclei has been suggested as the cause of correspondingly shorter and denser mitotic chromosomes. In another example, *Xenopus* chromosomes also start relatively long at the four-cell embryo stage, and they are shorter in the much smaller cells of a four-thousand-cell embryo towards the end of the cleavage division cycles (Zhou et al. 2023). In this case, the decreasing cytoplasm-to-nuclear ratio, and consequently the reducing quantity of available cytoplasmic condensin I complexes, has been proposed as the reason for shorter chromosomes.

Mitotic chromosome formation relies on the five-subunit protein complex condensin, that belongs to the Structural Maintenance of Chromosomes (SMC) family (Hirano 2016; Uhlmann 2016). Two types of condensin exist in vertebrates, including cytoplasmic condensin I, which gains access to chromosomes at the time of nuclear envelope breakdown in prophase. Condensin II, in turn, is a nuclear protein complex that becomes enriched on chromatin between prophase and telophase (Ono et al. 2003; Hirota et al. 2004; Ono et al. 2004; Gerlich et al. 2006). The relative concentrations of condensin I and II affect the chromosome length to width ratio by a yet unknown mechanism (Shintomi and Hirano 2011; Green et al. 2012). Other organisms, including budding and fission yeasts, contain only one type of condensin.

Condensin mediates mitotic chromosome formation by adding a layer of mitosis-specific long-range intra-chromosome interactions (Kakui et al. 2017; Gibcus et al. 2018; Kakui et al. 2022; Tang et al. 2023). The range of these intra-chromosome interactions differs between species. A wider spacing between detectable condensin binding sites correlates with farther-reaching interactions, as well as with wider chromosomes. Whether a causal relationship indeed links condensin spacing, chromatin interactions, and chromosome dimensions, remains to be investigated. The absolute numbers of condensin molecules also differs between species, with around twice as many condensin complexes present per fission yeast genome, compared to budding yeast (Breker et al. 2013; Carpy et al. 2014). The effect of quantitatively different condensin levels on chromosome dimensions also remains to be experimentally explored.

The dimensions of a mitotic chromosome arm furthermore depend on the fraction of the genome that is packed into each respective arm. Those chromosome arms that contain more DNA are longer, and they are also wider, an observation that holds true in both yeasts and vertebrates (Kakui et al. 2022). What is more, human chromosomes become progressively shorter and thicker the more time a cell spends in mitosis, and the width difference between short and long chromosome arms becomes increasingly prominent (Kakui et al. 2025). These chromosomes are therefore out-of-equilibrium structures, on the way to a steady state that is likely dictated by principles of polymer physics, but on biologically relevant timescales not typically reaching that state. In how far time spent in mitosis affects chromosome dimensions in other organisms has not yet been examined.

Here, we systematically explore chromosome size determinants in the fission yeast *Schizosaccharomyces pombe*. We study the effects of cell and nuclear size, of time spent in mitosis, and of the amount of available condensin. To measure fission yeast mitotic chromosome dimensions, we develop an unbiased line scan algorithm. Our analyses rule out previously suggested parameters, nuclear or cell size, or time in mitosis, as general chromosome size determinants. Instead, we find that condensin levels are a key determinant that defines chromosome dimensions in fission yeast.

## Results

### Chromosome size in fission yeast cells of increasing size

We set out by exploring whether the previously observed correlation between cell and nuclear size in interphase, and chromosome size in mitosis, is a general relationship that also applies to fission yeast. To address this question, we utilized a fission yeast strain harbouring an ATP analog sensitive *cdc2-asM17* allele (Aoi et al. 2014), allowing the use of the ATP analog 1NM-PP1 to inhibit the CDK cell cycle kinase and impose cell cycle arrest in G2 phase. During the G2 arrest, fission yeast cells continue to grow, with nuclear size increasing at the same rate (Neumann and Nurse 2007). We arrested cells in G2 for 3, 5 or 7 hours by 1NM-PP1 treatment, then released them to progress into mitosis where we blocked cell cycle progression again by transcriptional shut-off of the anaphase promoting complex co-activator Slp1 (Kakui et al. 2017). This protocol resulted in cells with increasingly larger nuclei entering mitosis. As a comparison, we arrested cells in metaphase by Slp1 shut-off without any prior G2 arrest.

We measured cell size from bright field images using image segmentation and cell masks generated by YeaZ (Dietler et al. 2020). Cell size in mitosis following 3 hours G2 arrest was comparable to the size of untreated cells in mitosis, while cell size in mitosis after 7 hours G2 arrest had more than doubled. To assess nuclear size, we mCherry-tagged the nuclear periphery protein Uch2 (Kouranti et al. 2010) and acquired fluorescent images in the TRITC channel, followed by image segmentation and nuclear mask generation using ilastik (Berg et al. 2019) (Fig. S1A). These analyses confirmed that, following G2 arrests of increasing durations, cells of increasing size entered mitosis with increasingly larger nuclei (Fig 1A and Fig S1B). Though we were unable to accurately measure nuclear size after 7 hours G2 arrest due to an increasingly abnormal nuclear morphology. We also measured the chromatin occupied area in interphase nuclei of increasing sizes, using ilastik to create image masks of the DNA binding dye 4′,6-diamidino-2-phenylindole (DAPI) stained chromatin area. This analysis revealed that chromatin spread out over much greater areas in larger nuclei. The expanded interphase chromatin substantially contracted again when cells with enlarged nuclei entered mitosis (Fig 1A).

**Fig 1.**
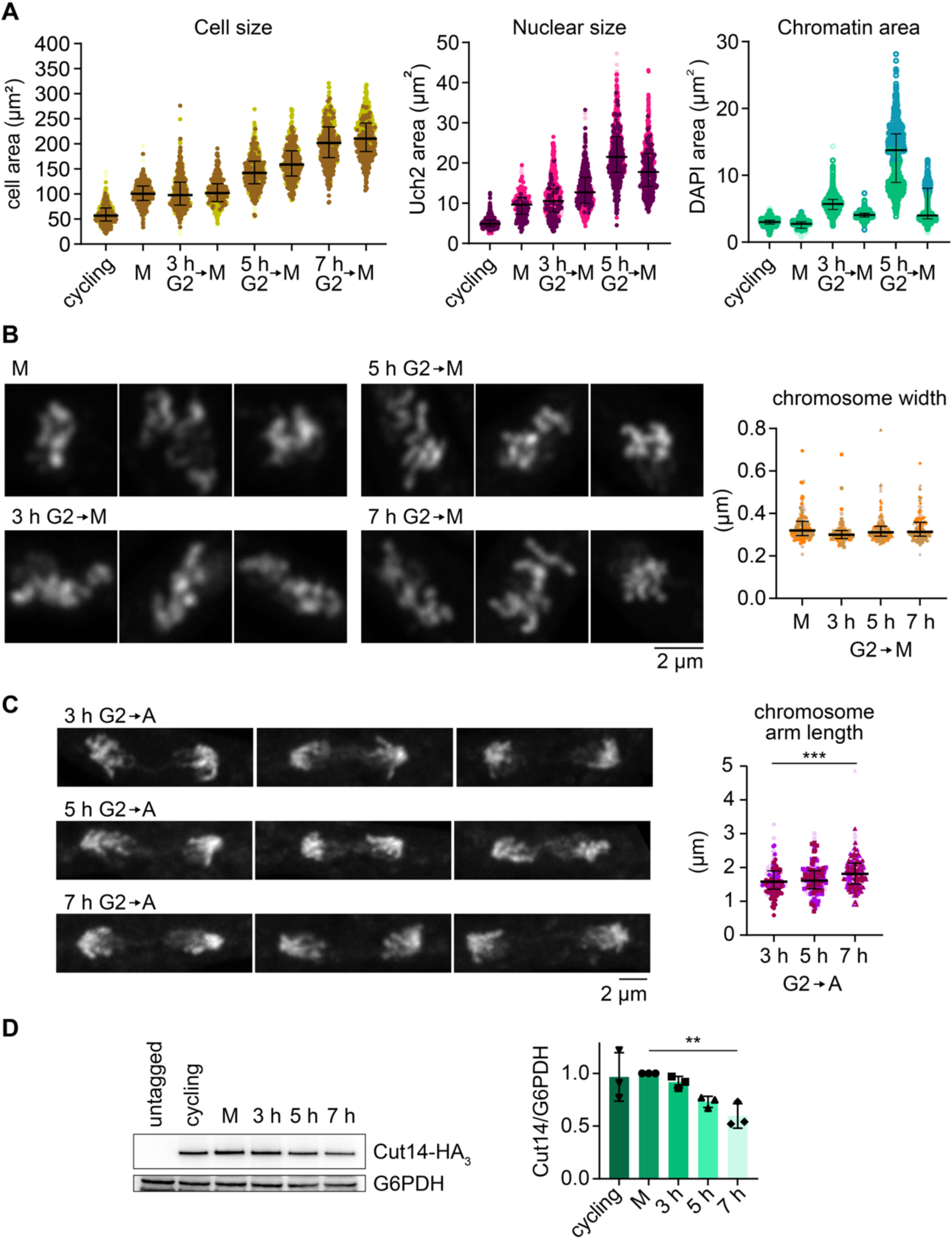
Chromosome size in larger cells. (**A**) Cell size, nuclear size and chromatin areas were determined in cells arrested for increasing lengths of time in G2, then released from G2 into mitotic arrest. Three biological repeat experiments were performed, colour coded, and aggregated. The medians and interquartile ranges of the aggregated measurements are indicated (Cell size: cycling, n = 615; M, n = 686; 3 h G2, n = 490; 3 h G2→M, n = 535; 5 h G2, n = 497; 5 h G2→M, n = 459; 7 h G2, n = 498; 7h G2→M, n = 507 – Nuclear size: cycling, n = 798; M, n = 327; 3 h G2, n = 796; 3 h G2→M, n = 945; 5 h G2, n = 940; 5 h G2→M, n = 659 – Chromatin area: cycling, n = 1087; M, n = 856; 3 h G2, n = 1129; 3 h G2→M, n = 1156; 5 h G2, n = 1079; 5 h G2→M, n = 902). (**B**) Representative Airyscan images of DAPI-stained chromosomes following G2 arrest for the indicated times and release into mitotic block, as well as measured mitotic chromosome widths. Three biological repeat experiments were performed, colour coded, and aggregated. The medians and interquartile ranges are indicated (M, n =115; 3 h G2→M, n = 82; 5 h G2→M, n = 163; 7 h G2→M, n = 154). (**C**) Representative Airyscan images of DAPI-stained chromosomes following G2 release at the indicated times into synchronous anaphase, as well as measured chromosome arm lengths. Three biological repeat experiments were performed, colour coded, and aggregated. The medians and interquartile ranges are indicated (3h G2→A, n = 142; 5 h G2→A, n = 152; 7 h G2→A, n = 225; *** *p* <0.0001, one-way ANOVA Tukey’s multiple comparisons test). (**D**) Representative immunoblot to analyse Cut14 levels at the indicated times, relative to G6PDH that served as the loading control. The Cut14/G6PDH ratios were quantified in three biological repeat experiments and normalized to the levels in the M population. Bars show the means and error bars the standard deviations (** *p* = 0.0070, one-way ANOVA Dunnett’s multiple comparisons test).

Following release from G2 arrest for 20 minutes into mitotic block due to Slp1 depletion, the majority of cells reached a mitotic state, indicated by the presence of short mitotic spindles (Fig S1C). At this time, we imaged DAPI-stained mitotic chromosomes using Airyscan superresolution microscopy (Huff 2015; Kakui et al. 2022) (Fig 1B). Z stacks of images were acquired and projected. We then used a semi-automated tool to measure the width of chromosome arms that we traced on these images (Fig S2A). The tool computationally straightens the traced chromosome arm, applies a sliding Gaussian fit along its length, and then records the average full width at half maximum intensity. This analysis revealed no significant changes to chromosome width, irrespective of the length of the G2 arrest, and consequently nuclear size, before entry into mitosis. The observation suggests that fission yeast mitotic chromosomes reach the same diameter irrespective of how widely distributed interphase chromatin was in nuclei of increasing sizes.

Next, we needed to measure chromosome arm lengths. While we could trace chromosome portions and determine their widths on the images of mitotically arrested cells, it was difficult to follow entire arms with enough certainty to establish their length. To measure chromosome arm lengths, we therefore changed our experimental approach. Instead of releasing G2 arrested cells into a mitotic block, we released cells to progress through mitosis and into anaphase, when we fixed and imaged cells as they segregated their chromosomes. At this time, chromosome arms are stretched out and their length could be measured by tracing on Airyscan images (Fig S2B). When we measured anaphase chromosome arm lengths following release from G2 arrests of increasing durations, we noticed gradual lengthening, which reached statistical significance when comparing cells after 3 and 7 hours of G2 arrest (Fig 1C). These results suggest that chromosome arms are longer, but not wider, in larger fission yeast cells.

Our above result is consistent with a scaling mechanism in which fission yeast cell size influences the length of its chromosome arms in mitosis. At the same time, we must consider possible confounding factors for this conclusion, stemming from differences between short and long cells. For example, chromosomes travel a farther distance during anaphase in longer cells, and the accompanying movement might have straightened chromosome arms more in longer than in shorter cells. Arm straightening in turn might have made chromosomes in larger cells appear longer in our measurements, when they might not in fact have been. Another confounding factor relates to the amount of available condensin in larger cells. When we analysed condensin levels in cells of increasing size by immunoblotting, we noticed that the relative condensin concentration decreased with increasing cell size, when compared to a metabolic housekeeping protein (G6PDH, Fig 1D). While the absolute number of condensin molecules is unlikely to decrease in larger cells, a decreasing condensin concentration in an enlarged cyto- and nucleoplasm might have nevertheless reduced the efficiency of its mitotic chromosome association. We will study the effect of an altered condensin concentration on chromosome size below. For now, to evaluate the merit of the concerns raised in this paragraph, we performed a reciprocal experiment using cells of a smaller size than wild type.

### Chromosome size in fission yeast cells of a smaller size

Given the confounding factors around our observation that chromosome arms are longer in larger fission yeast cells, we performed a reciprocal experiment. We measured chromosome size in fission yeast cells that were smaller than wild type. If chromosome size scales with cell size, we should expect chromosomes to be shorter in smaller cells. We employed two fission yeast cell size mutants in the cell cycle regulators Wee1 and PP2A. Cells lacking the CDK inhibitory kinase Wee1 (*wee1Δ*) or the major PP2A catalytic subunit (*ppa2Δ*) are smaller than wild type cells (Fantes and Nurse 1978; Kinoshita et al. 1993). Following 5.5 hours or 4.5 hours that were required, respectively, to arrest both types of cells in mitosis by Slp1 depletion, both mutant cells and their nuclei remained substantially smaller compared to their wild type counterparts (Fig 2A and Fig S3). We then measured chromosome width in mitotically arrested *wee1Δ* and *ppa2Δ* cells. Widths in *wee1Δ* cells did not significantly differ from chromosome widths in similarly mitotically arrested control cells, while widths in *ppa2Δ* cells were marginally reduced (Fig 2B).

**Fig 2.**
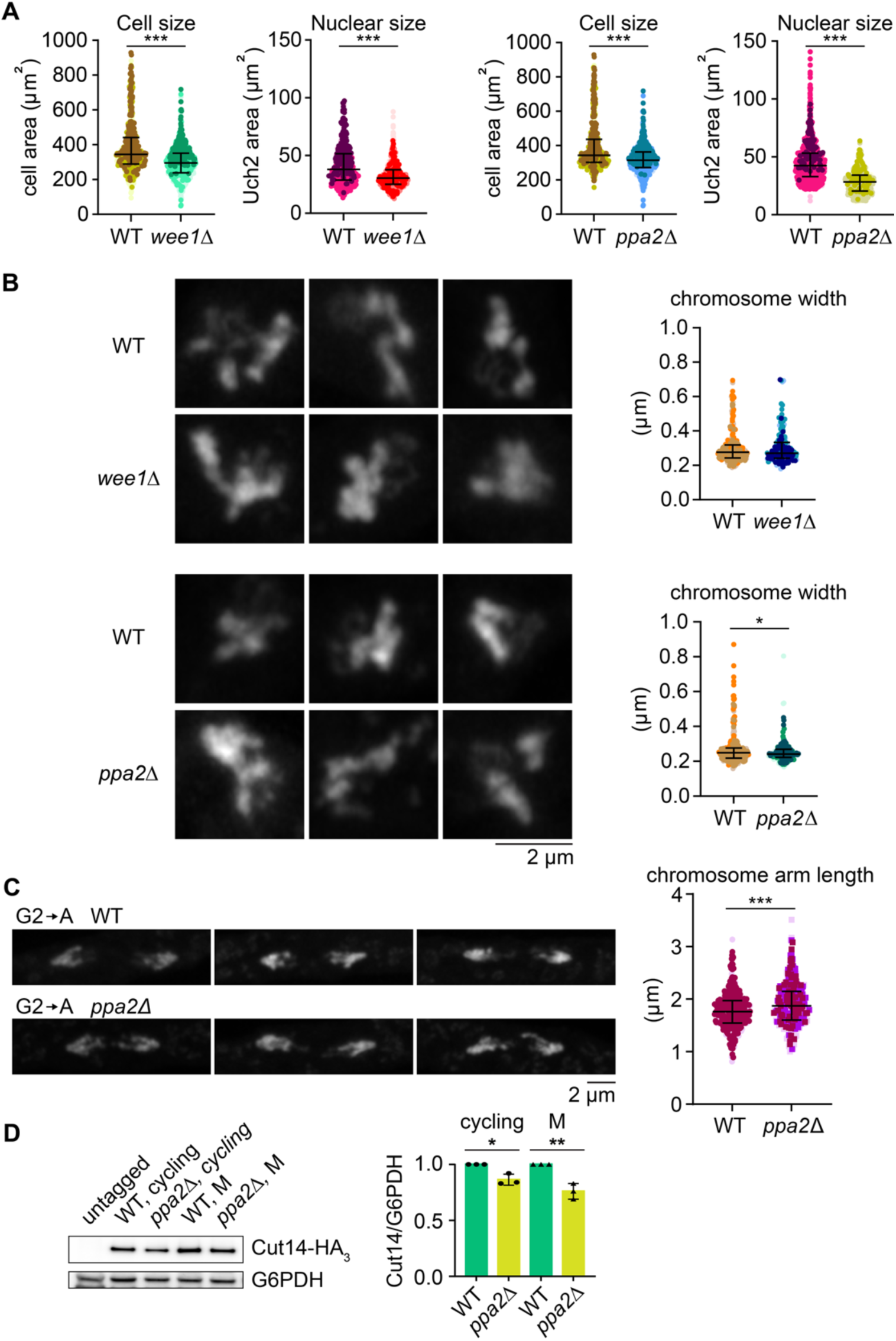
Chromosome size in smaller cells. (**A**) Cell size and nuclear size were determined in wild type (WT) and *wee1Δ* cells after 5.5 hours metaphase arrest. Three biological repeat experiments were performed, colour coded, and aggregated. The medians and interquartile ranges are indicated (Cell size: WT, n = 687; *wee1Δ*, n = 771; *** *p* <0.0001 – Nuclear size: WT, n = 418; *wee1Δ*, n = 422; *** *p* <0.0001, unpaired *t* tests). Similarly, WT and *ppa2Δ* cells were compared after 4.5 hours metaphase arrest (Cell size: WT, n = 613; *ppa2Δ*, n = 690; *** *p* <0.0001 – Nuclear size: WT, n = 954; *ppa2Δ*, n = 479; *** *p* <0.0001, unpaired *t* tests). (**B**) Representative Airyscan images of WT, *wee1Δ* and *ppa2Δ* DAPI-stained mitotic chromosomes at the respective arrest points, as well as measured chromosome widths. Three biological repeat experiments were performed, colour coded, and aggregated. The medians and interquartile ranges are indicated. (WT, n = 276; *wee1Δ,* n = 175; no significant difference – WT, n = 277; *ppa2Δ*, n = 193; * *p* = 0.0249, unpaired *t* tests). (**C**) Representative Airyscan images of DAPI-stained chromosomes following G2 release into synchronous anaphase of WT and *ppa2Δ* cells, as well as measured chromosome arm lengths. Three biological repeat experiments were performed, colour coded, and aggregated. The medians and interquartile ranges are indicated (WT, n = 422; *ppa2Δ*, n = 361; *** *p* <0.0001, unpaired t-test). (**D**) Representative immunoblot to analyse Cut14 levels in WT and *ppa2Δ* cells, asynchronously growing or arrested in G2, relative to G6PDH that served as the loading control. The Cut14/G6PDH levels were quantified in three biological repeat experiments and normalized to the levels found in WT cells. Bars show the means and error bars the standard deviations (* cycling, *p* = 0.0090; ** G2, *p* = 0.0003, one-way ANOVA Sidak’s multiple comparisons tests).

We next turned to measuring chromosome arm length during anaphase of small fission yeast cells following release from 3 hours cell synchronisation in G2 by CDK inhibition, a short G2 arrest time. We successfully obtained *ppa2Δ* cells containing the *cdc2-asM17* allele required for synchronisation by 1NM-PP1 treatment, but were unable to obtain *wee1Δ cdc2-asM17* cells. Against expectations, chromosome arms during anaphase of *ppa2Δ cdc2-asM17* cells were not shorter, but longer, when compared to the *cdc2-asM17* control (Fig 2C). This observation makes it unlikely that cell or nuclear size are directly linked to chromosome length. Rather, they suggest that another change that occurred in both larger and smaller fission yeast cells was the cause for anaphase chromosomes to appear longer.

As done for larger cells, we measured condensin levels in *ppa2Δ* cells by immunoblotting. This analysis revealed a reduced condensin concentration in *ppa2Δ* cells when compared to wild type cells, relative to the G6PDH housekeeping control protein (Fig 2D). While we do not know the reasons why condensin levels are reduced in these smaller cells, this observation opens the possibility that less condensin is available for mitotic chromosome formation in *ppa2Δ* cells compared to their wild type controls. The second possible confounding factor that we considered when looking at chromosome arm lengths, i.e. a farther separation distance during anaphase in longer cells, is of course not applicable in these shorter cells. Taken together, we find no consistent correlation between fission yeast cell and nuclear size in interphase, and chromosome size in mitosis. Instead, we find that reduced relative condensin levels in both larger and smaller fission yeast cells correlate with somewhat longer chromosome arms.

### Condensin-depletion causes wider and longer chromosomes

To test whether reduced condensin levels might be an underlying reason for longer chromosome arms in both larger and smaller fission yeast cells, we directly altered available condensin levels by tagging the condensin subunit Cut14 with an auxin-inducible degron (AID) tag (Kanke et al. 2011; Kakui et al. 2017). Titrating the auxin concentration that we added to the culture medium allowed gradual condensin depletion (Fig 3A). At the same time as adding auxin, we supplemented the culture with thiamine to shut off Slp1 transcription and thereby induce mitotic arrest (Fig S4A). After five hours, mitotic chromosomes that formed in the presence of decreasing condensin concentrations were imaged by Airyscan microscopy. Even a small (approximately 30%) reduction of condensin levels caused by the addition of 25 µM auxin resulted in a measurable and significant widening of the resulting chromosomes, an effect that increased at 50 µM auxin when Cut14 levels were less than half that of wild type (Fig 3B). When condensin was further depleted (500 µM auxin) no measurable chromosomes could be seen in the mitotic arrest. These results suggest that condensin is a limiting component that determines chromosome width.

**Fig 3.**
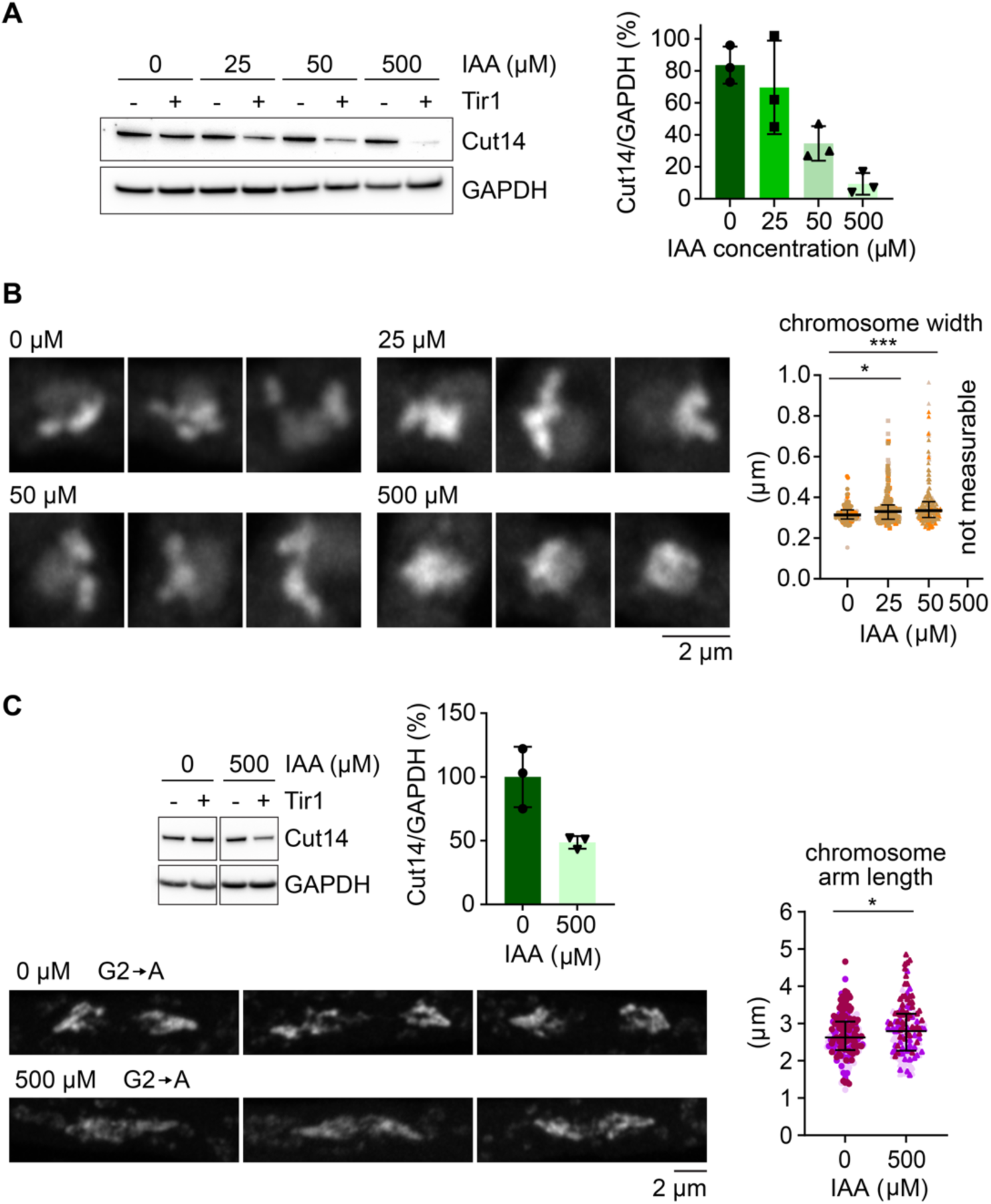
Chromosome size following condensin titration. (**A**) Representative immunoblot to analyse Cut14 levels during its depletion using the indicated auxin concentrations in Cut14-aid strains expressing, or not, Tir1. GAPDH served as loading control. The Cut14/GAPDH ratios were quantified in three biological repeat experiments and normalised to the levels observed in the strain lacking Tir1. Bars show the means and error bars the standard deviations. (**B**) Representative Airyscan images of DAPI-stained mitotic chromosomes in cells treated with the indicated auxin concentrations, as well as measured mitotic chromosome widths. Three biological repeat experiments were performed, colour coded, and aggregated. The medians and interquartile ranges are indicated. (0 µM, n = 100; 25 µM, n = 158; 50 µM, n = 183; * *p* = 0.0296, *** *p* = 0.000, one-way ANOVA Tukey’s multiple comparisons tests). (**C**) Representative Airyscan images of DAPI-stained chromosomes following G2 release into synchronous anaphase of Cut14-aid cells treated with 0 or 500 µM auxin, as well as measured chromosome arm lengths. Three biological repeat experiments were performed, colour coded, and aggregated. The medians and interquartile ranges are indicated (0 µM, n = 209; 500 µM, n = 144; * *p* = 0.0177, unpaired *t* test). A representative immunoblot to analyse Cut14 depletion in this experimental setting is shown, as well as quantification relative to GAPDH that served as a loading control in triplicate biological repeats. Bars show the means and error bars the standard deviations.

Next, we assessed the impact of condensin levels on chromosome length. We used our G2 synchronisation protocol and, at the same time as 1NM-PP1, we added auxin to the growth medium. To avoid complications from cell elongation, we again limited the G2 synchronisation period to 3 hours, after which we released cells for progression into anaphase. Using this depletion protocol, 500 µM auxin addition resulted in an approximate halving of condensin levels. As the consequence, anaphase chromosome arms became significantly longer (Fig 3C). These observations suggest that condensin levels control chromosome dimensions, with decreasing condensin levels resulting in both wider and longer chromosomes.

### Condensin and nuclear size in interphase

It has been suggested that condensin constrains interphase chromatin, thereby limiting its entropic expansion and thereby controls nuclear size in both *Drosophila* and human cells (George et al. 2014). To examine if such a role of condensin is conserved in fission yeast, we measured cell and nuclear size following condensin depletion (Fig S4B). Against expectations, we did not see larger nuclei in condensin-depleted cells. On the contrary, we found that nuclei were smaller in mitotically arrested cells with reduced condensin levels. Cell size in mitosis was also smaller in condensin-depleted cells. We do not know the reason for reduced cell and nuclear size following condensin reduction, but cannot exclude that a role of condensin in transcriptional regulation might have contributed (Lancaster et al. 2021). We conclude that nuclear size, at least in fission yeast, is not limited by a role of condensin in constraining the interphase chromatin volume. Rather, we have seen above that it is the nuclear size that limits how far interphase chromatin can spread (Fig 1A).

### Extra condensin causes increased chromosome compaction

Chromosomes were wider and longer upon condensin depletion, consistent with the possibility that condensin levels regulate chromosome dimensions in fission yeast. However, larger chromosomes at reduced condensin levels could merely be an unphysiological intermediate on the way to chromosomes losing all their shape without condensin (Fig. 3B) (Saka et al. 1994; Sutani et al. 1999). If condensin levels were indeed a limiting factor that determines fission yeast chromosome size, then increasing the available condensin should result in smaller chromosomes. To investigate this possibility, we introduced an additional 2^nd^ copy of the genes encoding each of the five condensin subunits, Cut3, Cut14, Cnd1, Cnd2 and Cnd3, under control of their native upstream and downstream regulatory regions, at ectopic loci in the fission yeast genome (referred to as the “2^nd^ copy” strain). The additional copies of each condensin gene resulted in a detectable condensin level increase, as seen by immunoblotting (Fig 4A). Measuring chromosome dimension in mitotically arrested cells revealed significantly thinner chromosomes in the 2^nd^ copy strain, as compared to the wild type control (Fig 4B). Similarly, measuring chromosome arm length following release of G2 synchronised cells into anaphase revealed that chromosome arms were significantly shorter due to the increased condensin dosage (Fig 4C). These observations demonstrate that condensin levels indeed limit chromosome compaction during fission yeast mitosis. The greater the condensin levels the more compact, shorter and thinner, chromosomes become.

**Fig. 4.**
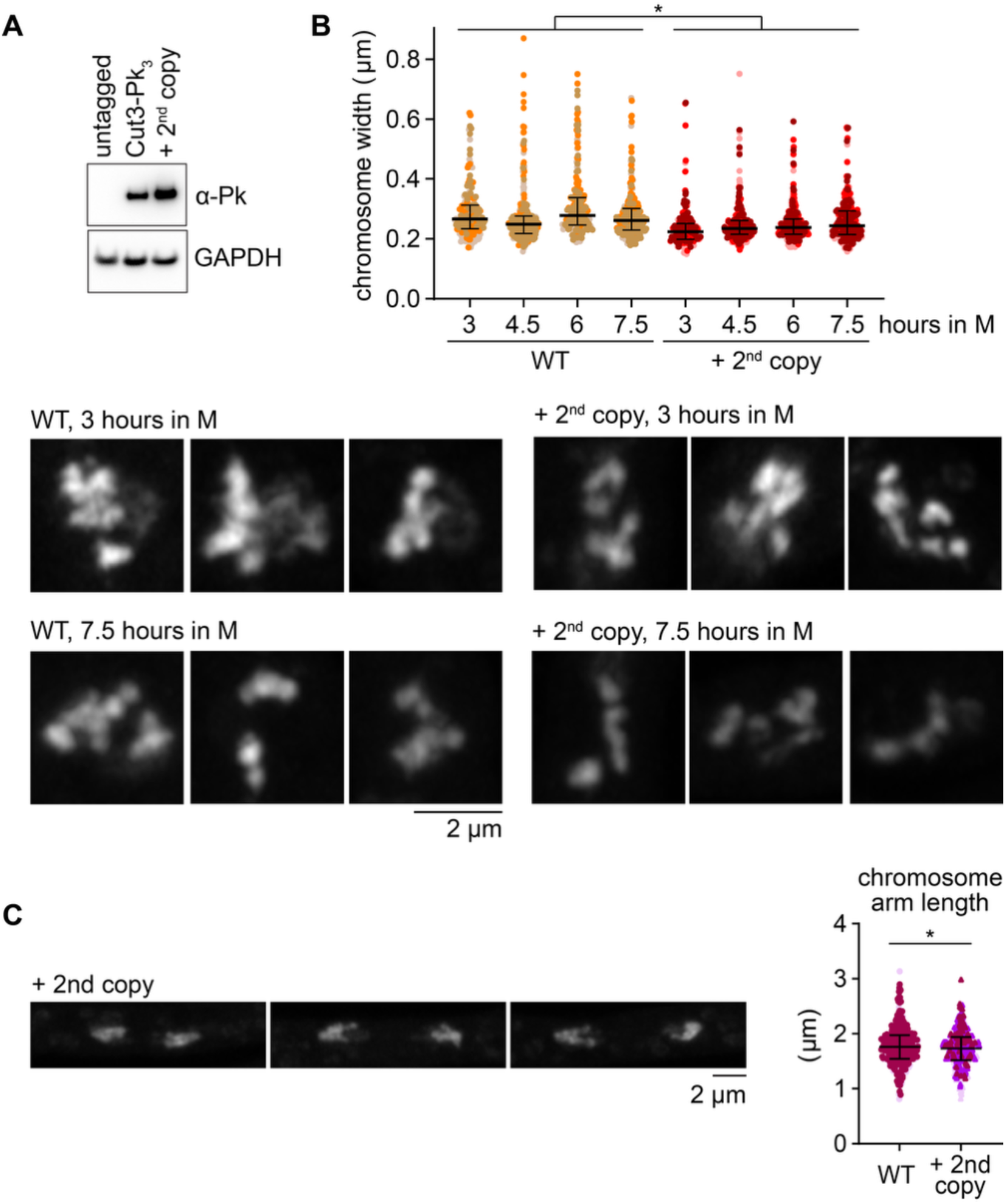
Chromosome size at increased condensin dosage. **(A)** Immunoblot comparing endogenous Cut3-Pk_3_ levels to those in the 2^nd^ copy strain. An untagged Cut3 strain was included as control. GAPDH served as the loading control. (**B**) Representative Airyscan images of DAPI-stained mitotic chromosomes in WT and 2^nd^ copy strains during the indicated times in a metaphase arrest, as well as measured chromosome widths at these and additional times. Three biological repeat experiments were performed, colour coded, and aggregated. The medians and interquartile ranges are indicated. (WT: 3 hours, n = 176; 4.5 hours, n = 278; 6 hours, n = 207; 7.5 hours, n = 242 – 2^nd^ copy: 3 hours, n = 208; 4.5 hours, n = 273; 6 hours, n = 284; 7.5 hours, n = 272; time effect: not significant, * genotype effect: *p* = 0.0353, two-way ANOVA). (**C**) Representative Airyscan images of DAPI-stained anaphase chromosomes in the 2^nd^ copy strain, as well as measured chromosome arm lengths, compared to arm lengths in a WT strain. Three biological repeat experiments were performed, colour coded, and aggregated. The medians and interquartile ranges are indicated (WT, n = 422; 2^nd^ copy n = 343; * *p* = 0.0304, unpaired *t* test).

### Time in mitosis does not affect fission yeast chromosome dimensions

Vertebrate chromosomes have been observed to undergo continuous shape changes throughout the course of mitosis, with chromosomes become gradually shorter and thicker over time (Kakui et al. 2025). Quantification of these shape changes showed that mitotic chromosomes are out of equilibrium structures on the way to a steady state that is reached before anaphase onset only by the shortest, but not by longer chromosome arms. We therefore investigated whether fission yeast chromosomes similarly change shape in cells that are arrested in mitosis. To do so, we kept both a wild type strain, as well as the 2^nd^ copy strain, in a mitotic arrest due to Slp1 shut-off for an extended period from 3 hours up to 7.5 hours (Fig S4C). However, when we measured chromosome widths at successive times, we did not detect any measurable change over time (Fig 4B). Condensin levels remained constant during the time of the arrest (Fig S4D). These observations suggest that, unlike human chromosomes, fission yeast chromosomes reach a steady state faster, maybe due to their smaller size.

## Discussion

Here, we performed a systematic study in the search for chromosome size determinants in fission yeast. To our surprise, we did not find a consistent correlation between cell and nuclear size in interphase and chromosome size in mitosis. While chromatin was spread out over a far greater area in the larger nuclei of larger interphase cells, the ensuing mitotic chromosomes presented a width that was indistinguishable from wild type. Though in both larger and in smaller cells, we found that chromosome arms were measurably longer. This feature correlated with a reduced condensin concentration in both larger and smaller cells. Reduced condensin levels in large cells could stem from a dilution effect, where the absolute condensin amount remains constant but is diluted in a growing cellular environment. In contrast, it is not immediately obvious why condensin levels were reduced in small cells. Condensin was reported to interact with PP2A in vertebrates (Takemoto et al. 2009). Whether a condensin– PP2A interaction exists in fission yeast that might have impacted on condensin levels or function in smaller cells, or cells lacking Ppa2, remains to be explored. How the levels of chromosomal proteins scale with cell size remains a topic of wider interest (Schwaffer et al. 2021).

If longer chromosomes in both larger and smaller fission yeast were caused by reduced condensin levels, why did we not observe that chromosomes were also wider, as we did when we experimentally depleted condensin? On the contrary, chromosomes in *ppa2Δ* cells appeared somewhat thinner. It is possible that small changes to chromosome length are more reliably discernible than small alterations in chromosome width. Alternatively, factors additional to reduced condensin levels could have contributed to the chromosome size changes that we observed in both larger and smaller fission yeast. We note that reduced condensin I levels were put forward as the reason for shorter chromosomes during *Xenopus* development (Zhou et al. 2023), the opposite of the tendency that we observed in fission yeast. A causal relationship between condensin levels and chromosome size was not tested as part of the *Xenopus* experiments. In the future, a quantitative assessment of chromosome-bound condensin, though experimentally challenging, should clarify whether less condensin is found on chromosomes in large and small fission yeast cells, and is the reason for the observed chromosome size changes.

Unlike in vertebrates, where chromosomes keep changing shape for as long as it possible to observe them (Mora-Bermúdez et al. 2006; Shintomi et al. 2017; Gibcus et al. 2018; Kakui et al. 2025), fission yeast chromosomes reached an apparently constant appearance in a relatively short time. Fission yeast condensin might work in ways different from vertebrate condensin, though we have no indication of overt molecular or structural differences. More likely, therefore, their smaller size allows fission yeast chromosomes to reach a steady state faster. It will be interesting to establish the dimensions of this state, though the limitations of optical microscopy that we have encountered in our study make it hard to directly measure both the length and width of individual fission yeast chromosome arms. At first glance, the final fission yeast chromosome arm shape appears elongated.

The biggest impact on fission yeast chromosome size that we could establish came from the cellular condensin levels. The more condensin the shorter and thinner the chromosomes became. What can we learn from this behaviour about the mechanism by which chromosomes form? The wider cellular context exerted little influence on what chromosomes eventually look like. Instead, chromosome formation largely appears autonomous, depending on chromatin and the action of the condensin complex. Two prominent models for chromosome formation by condensin have been considered in the recent literature, the loop extrusion and the loop capture models (Kinoshita et al. 2022; Kim et al. 2023; Tang et al. 2023; Uhlmann 2025). The loop extrusion model foresees that more condensin results in a greater number of loops along chromosomes, resulting in smaller loops and consequently thinner and also longer, chromosomes (Goloborodko et al. 2016). This prediction was not met by our experimental observations. Increased condensin levels did make chromosomes thinner, but also shorter. The loop capture scenario in turn might accommodate the observation of both thinner and shorter chromosomes. If loop capture opportunities remain unsaturated, such that additional condensin creates additional loops, then these additional loops could result in further chromosome compaction in both length and in width (Kakui et al. 2025). In the future, measuring condensin’s loop architecture, in dependence on the condensin concentration, will yield further insight into the mechanisms by which chromosomes take shape.

## Materials and Methods

### *S. pombe* strains and culture

All strains used in this study are listed in Table S1. Strains were constructed using PCR-based gene targeting, and by genetic crossing and tetrad dissection. To construct the Cut14-aid strain, the *cut14*^+^ gene was fused to the sequence encoding the auxin-inducible IAA17 degron module in a strain harbouring Skp1-TIR1 (Kakui et al. 2020). To obtain the 2^nd^ copy condensin strain, three plasmids were created harbouring ectopic copies of the five condensin subunits (detailed in Table S2). These plasmids were linearised for integration at their respective marker loci in a host strain expressing Cut3-Pk_3_ also from the endogenous locus.

To achieve metaphase arrest, strains harbouring the kan^R^::P*_nmt_41-slp1*^+^ allele (Kakui et al. 2017) were cultured in Edinburgh minimal medium (EMM) supplemented with 2% glucose and 3.75 mg/ml glutamic acid, adenine, leucine, uracil and histidine supplements at 25°C overnight. The cells were filtered and resuspended in yeast extract (YE) medium containing 3% glucose, four amino acids supplements (YE4S), and 5 µg/ml thiamine to shut off Slp1 expression for the indicated times at 25°C. To induce Cut14-depletion in the kan^R^::P*_nmt_41-slp1*^+^ background, the Cut14-aid strain was cultured as above, with auxin supplement at indicated concentrations at the time of shifting to thiamine-containing YE4S medium for five hours at 25°C. To arrest cells in G2, strains harbouring the *cdc2-asM17* allele (Aoi et al. 2014) were cultured in EMM4S medium at 30°C overnight. 1 µM 1-NMPP1 and 5 µg/ml thiamine were added the next day for 3, 5, or 7 hours. To transition from G2 to a mitotic arrest, cells were filtered and washed at the indicated times and released into thiamine-containing YE4S medium for 20 minutes at 30 °C. To release cells from G2 to anaphase, the *cdc2-asM17* strains were treated as described above, except for omitting thiamine addition to the EMM4S or YE4S media. Samples were collected at 2-minute intervals.

### Yeast chromosome width measurements

Collected cells were fixed with 70% ethanol. After fixation, the samples were resuspended in 10% glycerol containing 0.25 µg/ml DAPI. Mitotic chromosomes were imaged by Airyscan microscopy (Huff 2015) using a 63x/1.40 NA objective lens. Images were acquired along the z-axis with the first/last option and optimal spacing. Snapshots of mitotic spindles that span across the nucleus were taken to monitor the mitotic cell population. To measure chromosome width, chromosome arms were segmented using a custom Fiji macro (Schindelin et al. 2012). We selected chromosomes by placing points along the long chromosome axis on a maximum intensity projection of a 3D image stack using the “polyline” tool. The line selection along the chromosome axis was straightened using the “Straighten” tool in Fiji. The image was then cropped along the axis of the straightened chromosome and the region of interest saved. The ROI was overlaid on the original image to ensure that each chromosome was selected only once.

Image analysis of the segmented chromosome arms was performed in MATLAB. Chromosome images were first aligned so that the chromosome long axis is aligned with the vertical axis of the image. For each chromosome, the image is thresholded and a binary mask is created. Holes in the mask are filled, and a convex hull of the mask is generated. At this point, the mean width of the binary mask rows is measured, and any individual row that has a width less than 70% of the mean is removed. This step removes any telomere ends from the dataset. The first rows from each chromosome end where the width is greater than 70% of the mean are marked as *border rows.* The binary mask is then expanded by 2 pixels, and applied to the chromosome image, setting all background intensities to zero.

The image was then normalised, and a Gaussian fit applied to the image intensity in each row along the chromosome axis. The rows are then filtered, any row with a r^2^ less than the specified threshold, or with a confidence interval that is an outlier as measured by the *rmoutliers* function, were removed from the dataset. The full width half maximum (FWHM) value of each Gaussian was then found and compared to the all the FWHM for each row in the chromosome image. Any row with a FWHM deemed an outlier by the *rmoutliers* function is removed from the dataset. The remaining rows are then combined to give a mean measurement of the width for the chromosome, as well as standard deviation and standard error of the mean.

The FWHM and standard error of the mean are plotted for each chromosome dataset (i.e. multiple chromosomes under the same condition). A histogram of the FWHM is also generated per dataset. A montage image for inspection is also created, where the rows that are measured are overlaid in blue on the original image, with the border rows in red. The chromosome width measurements were done in triplicate biological repeat experiments, at least 50 nuclei were imaged for each experiment.

All custom image analysis tools in Fiji and MATLAB, with usage instructions, have been deposited with the Zenodo repository, where they can be accessed with the following DOI: 10.5281/zenodo.15657164

### Cell size, nuclear size and chromatin area measurements

Collected cells were fixed with 70% ethanol or 3.6% formaldehyde. Images of bright field, TRITC (visualising Uch2-mCherry), DAPI and FITC (visualising GFP-Atb2) channels were acquired with a DeltaVision or Zeiss Observer equipped with a 63x objective lens. At least 200 cells were imaged in 7 slices at 0.5 µm spacing along the z-axis for monitoring cell area, nuclear area, chromosome area and the population displaying mitotic spindles, respectively. Corresponding cell masks were created using YeaZ (Dietler et al. 2020), while the nuclear and chromatin area masks were generated by ilastik (Berg et al. 2019). The masks were then inspected using Fiji. Objects that were not successfully masked, or did not display a mitotic spindle, were removed using the wand tool. Conversely, when measuring nuclear or chromatin areas in cycling cell populations, cells in metaphase that showed mitotic spindles were removed from the analysis. Subsequently, the look up table was converted to a colour range for thresholding, and the scale was converted from pixels to µm. The area was then measured by selecting the ‘analyse particles’ option. Triplicate biological repeat experiments were performed in all cases.

## Acknowledgments

We would like to thank Ying Gu, Thomas Hammond, Yasu Kakui, Sofi Mebrate, Snezhana Oliferenko, Matt Renshaw, Billy Whyte and Theresa Zeisner for reagents and advice, Rocco D’Antuono and Matt Renshaw from the Crick Advanced Light Microscopy STP and Gavin Kelly from the Bioinformatics & Biostatistics STP for help with microscopy and statistics, as well as all our laboratory members for discussions and comments on the manuscript. This work was supported by a Wellcome Trust Investigator Award (220244/Z/20/Z) and the Francis Crick Institute that receives its core funding from Cancer Research UK, the UKRI Medical Research Council, and the Wellcome Trust (cc2137).

## Author contributions

P.-S.W. and F.U. conceived the study, P.-S.W. performed all experiments, T.F. developed the image analysis tools, P.-S.W. and F.U. wrote the manuscript with input from T.F.

---

**Fig S1.**
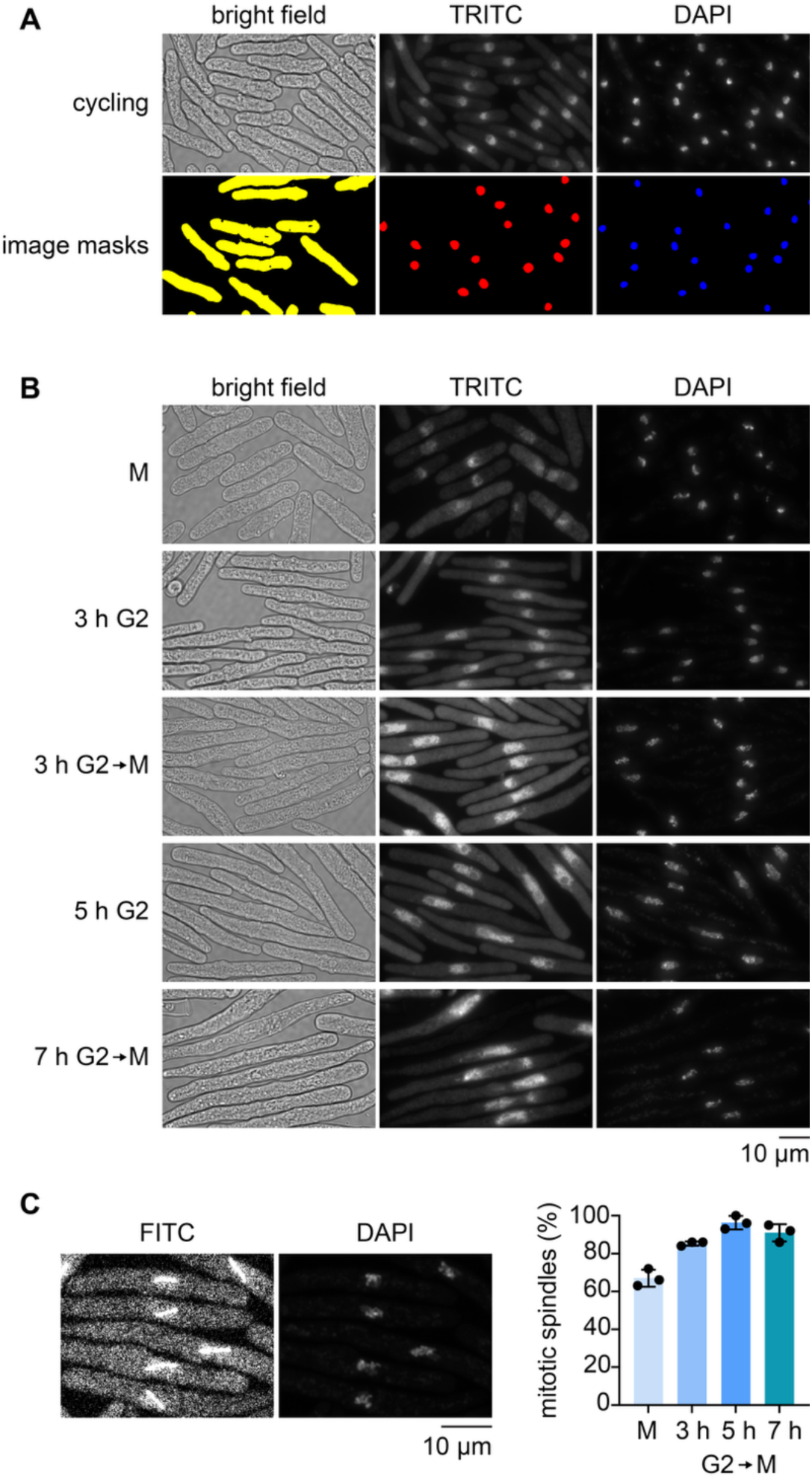
Chromosome size in larger cells – supporting information. (**A**) Representative bright field, TRITC (visualising Uch2-mCherry), and DAPI images of a cycling cell population. Image masks of the cell outlines generated by YeaZ, and nuclear and chromatin masks generated by ilastik are shown alongside. (**B**) Representative bright field, TRITC (visualising Uch2-mCherry), and DAPI images of cells following the indicated lengths of G2 arrest, released or not for an additional 20 minutes into a mitotic block. (**C**) Representative FITC (visualising GFP-Atb2) and DAPI images of 3 h G2→M cells. The fraction of cells displaying mitotic spindles was counted following the indicated times in G2 arrest and release into mitotic block, in each of the three biological repeat experiments reported in Figures 1A and B. Bars show the means and error bars the standard deviations.

**Fig S2.**
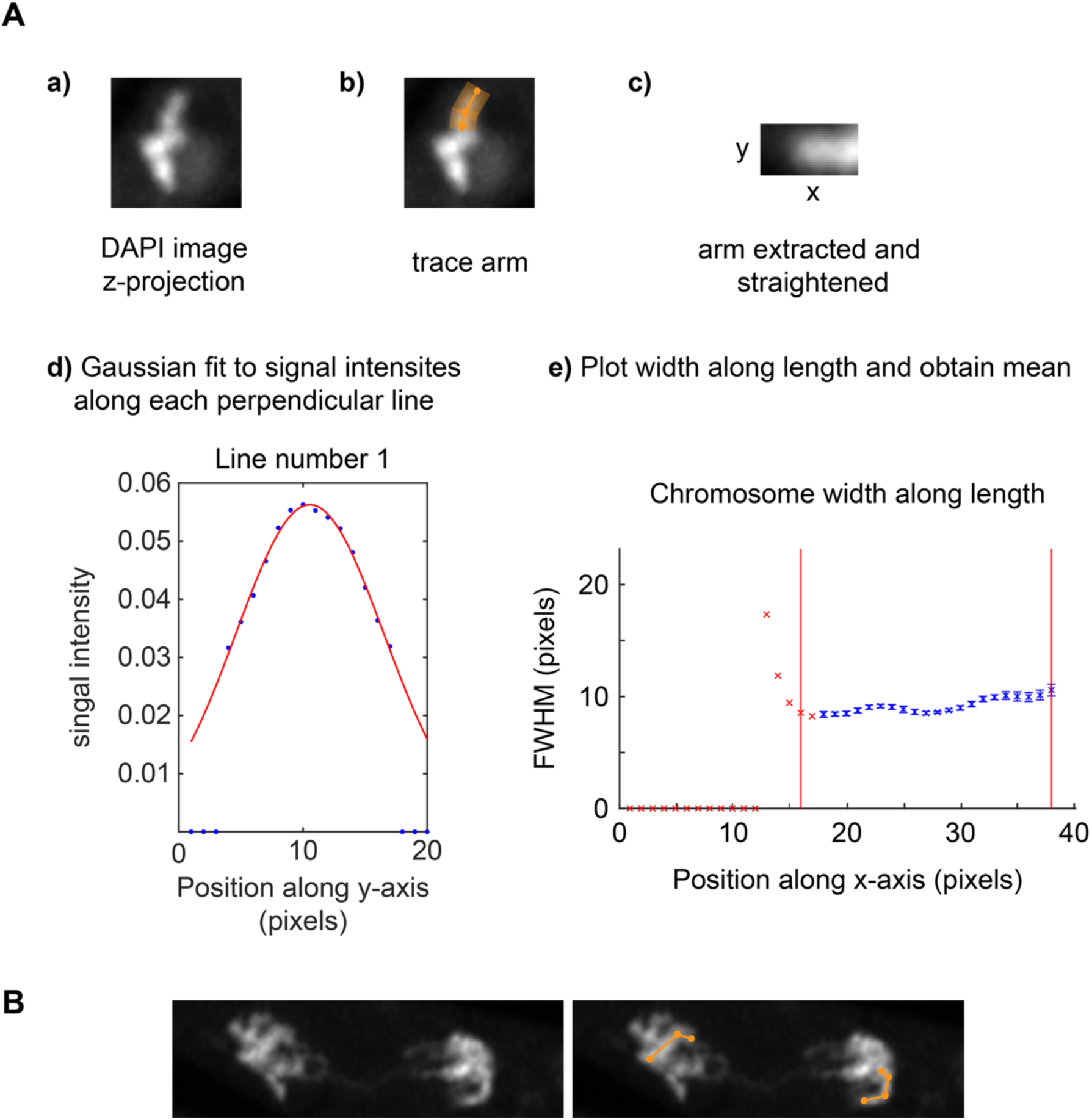
Methods to measure chromosome widths and lengths. (**A**) Example mitotic chromosome image and the steps that were followed to measure the average full width at half maximum (FWHM) of a visible continuous chromosome body. See Materials and Methods for details. (**B**) Illustration of a chromosome arm length measurement. The segregating arms were traced with the orange segmented lines. The corresponding arm lengths were then recorded using the analysis option in Fiji. All arms pointing in the direction of anaphase separation were included in the measurements.

**Fig S3.**
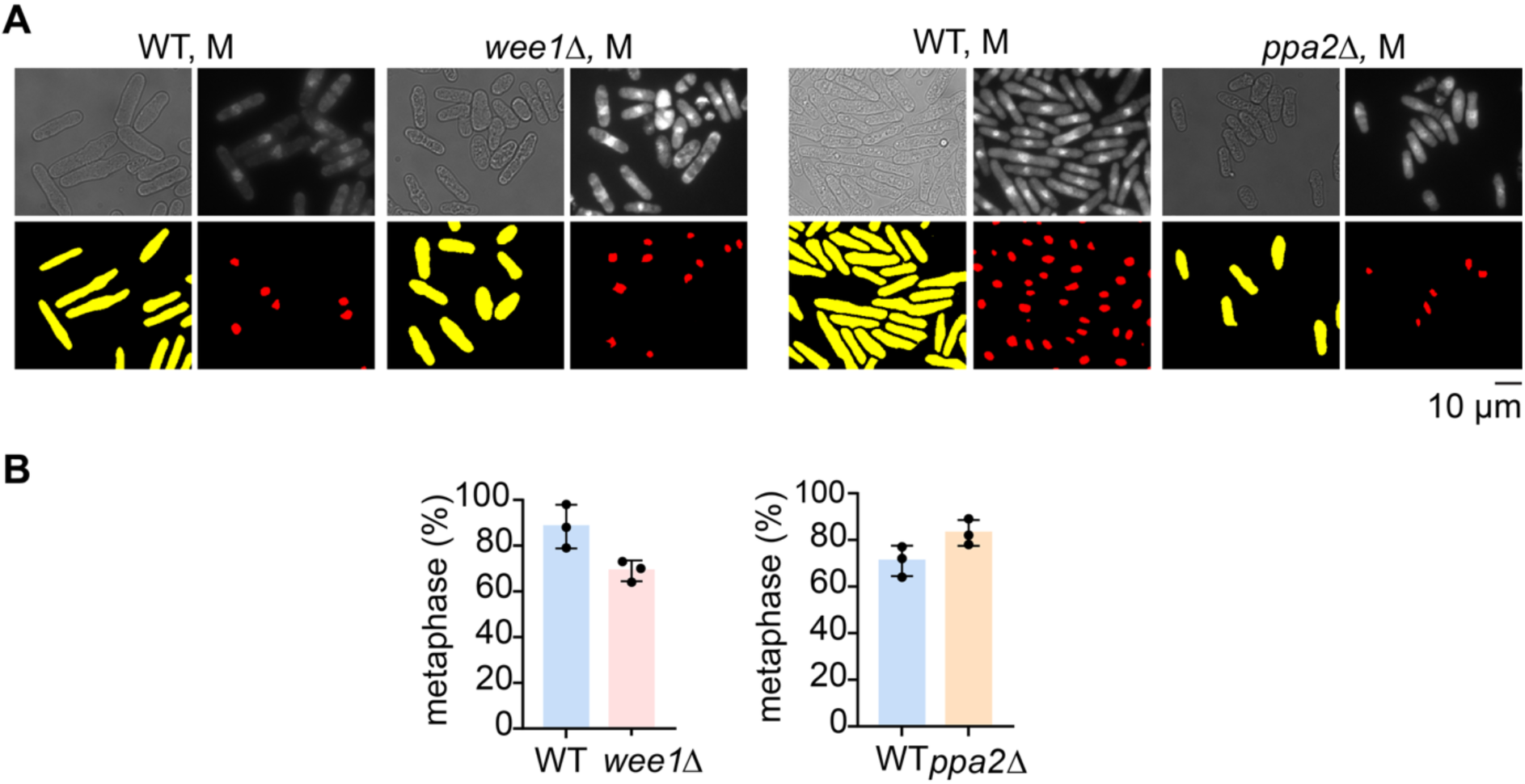
Chromosome size in smaller cells – supporting information. (**A**) Representative bright field and TRITC (visualising Uch2-mCherry) images of WT, *ppa2Δ*, and *wee1Δ* cells following 4.5 h and 5.5 h of mitotic arrest, respectively. The corresponding cell masks and nuclear masks generated by YeaZ and ilastik are shown alongside. (**B**) Percentage of WT and *ppa2Δ* cells that presented mitotic spindles after 4.5 hours, as well as percentage of WT and *wee1Δ* cells that showed mitotic spindles after 5.5 hours in a metaphase arrest. 100 cells were scored in each of the three biological replicate experiments shown in Figures 2A-C.

**Fig S4.**
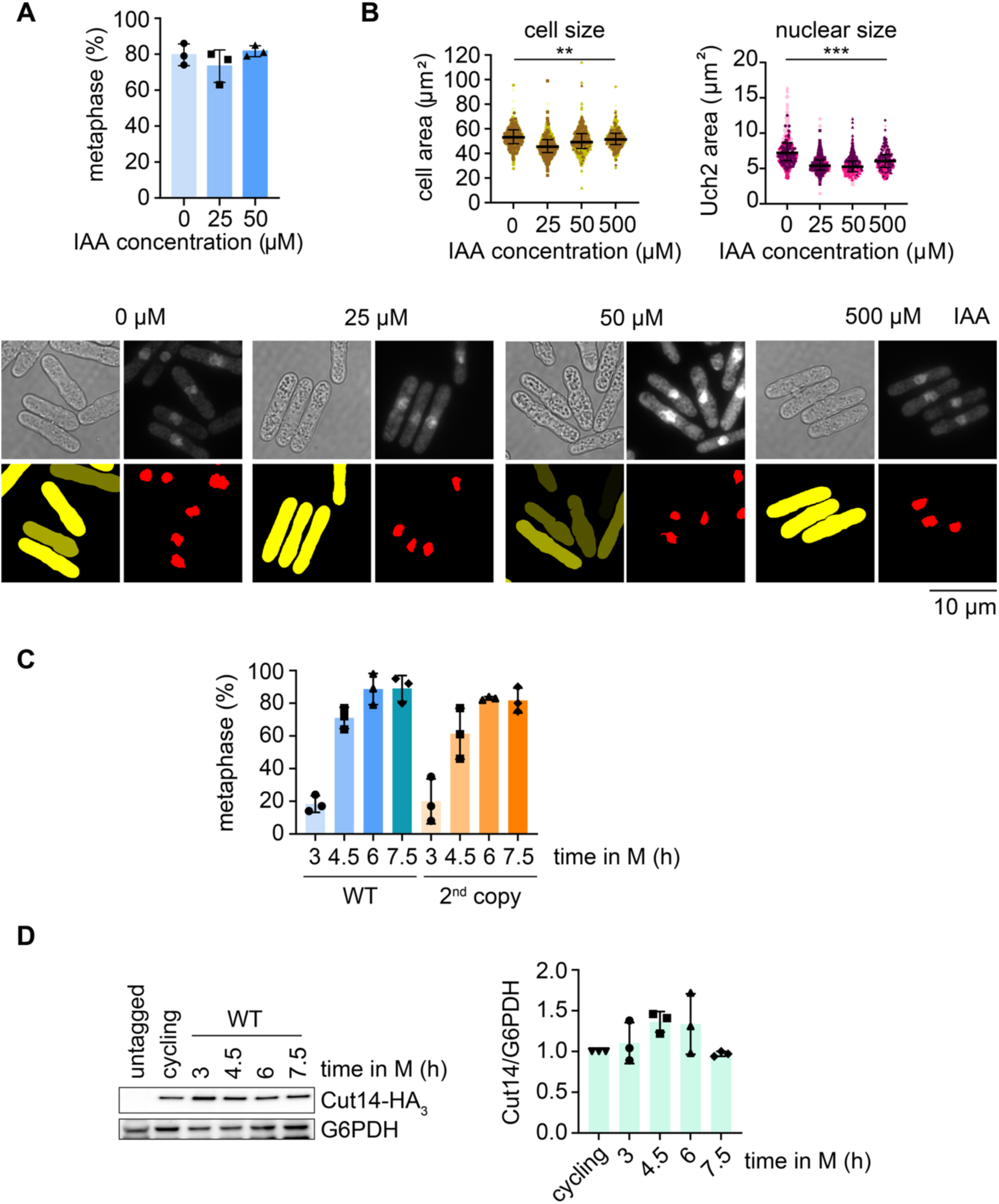
Condensin dosage – supporting information. (**A**) The percentage of cells displaying mitotic spindles in the experiment shown in Figs 3A, B at the indicated auxin concentrations. 100 cells were scored from each of the three biological repeat experiments. Bars show the means and error bars the standard deviations. (**B**) Representative bright field and TRITC (visualising Uch2-mCherry) images, showing cells treated with the indicated auxin concentrations, and their nuclei, following five hours in mitotic arrest. The corresponding cell masks and nuclear masks generated by YeaZ and ilastik are shown alongside. Cell and nuclear areas were measured in three biological repeat experiments, colour coded, and aggregated. The medians and interquartile ranges are indicated (Cell size: 0 µM, n = 711; 25 µM, n = 865; 50 µM, n = 793; 500 µM, n = 685; ** *p* = 0.0002 – Nuclear size: 0 µM, n = 465; 25 µM, n = 563; 50 µM, n = 513; 500 µM, n = 412; *** *p* < 0.0001, one-way ANOVA, Dunnett’s multiple comparisons tests). (**C**) Percentage of WT and 2^nd^ copy cells that displayed mitotic spindles during the mitotic arrest timecourse experiment in Fig 4B. 100 cells were scored in each of the three biological repeat experiments. Bars show the means and error bars the standard deviations. (**D**) Cut14 protein levels in WT cells were followed over the indicated times in a mitotic arrest. G6PDH served as the loading control. A representative immunoblot is shown, as well as quantification of the Cut14/G6PDH ratios in the three biological repeat experiments shown in Fig 4B, normalised to the ratio seen in asynchronously growing (cycling) cells. Bars show the means and error bars the standard deviations.

**Table S1.** Yeast strains used in this study.

| Strain | Genotype | Figure |
| --- | --- | --- |
| 7009 | *h<sup>-</sup> Kan::P<sub>nmt41</sub>-slp1<sup>+</sup> SV40 P-GFP-atb2::LEU2*<br>*Uch2-mCherry::ura4<sup>+</sup> cut14-IAA17-ura4<sup>+</sup>*<br>*ade6::ade6<sup>+</sup>-Padh15-skp1-OsTIR1-natMX6-*<br>*Padh15-skp1-AtTIR1-2NLS* | 3A, 3B, S4A-S4B |
| 7011 | *h<sup>-</sup> Kan::P<sub>nmt41</sub>-slp1<sup>+</sup> SV40 P-GFP-atb2::LEU2*<br>*Uch2-mCherry::ura4<sup>+</sup> cut14-IAA17-ura4<sup>+</sup>* | 3A, 3B |
| 6963 | *h<sup>-</sup> Kan::P<sub>nmt41</sub>-slp1<sup>+</sup> SV40 P-GFP-atb2::LEU2*<br>*Uch2-mCherry::ura4<sup>+</sup> leu1-32* | 4B, 4C, S4D |
| 7054 | *h<sup>-</sup> kan::P<sub>nmt41</sub>-slp1<sup>+</sup> Cut3-3Pk::HPH*<br>*ura4<sup>+</sup>::cut3-3Pk,cut14-natMX*<br>*lys3<sup>+</sup>::cnd3-BsdMX his5<sup>+</sup>::cnd2-3XHA,cnd1-bleMX*<br>*SV40 P-GFP-atb2::LEU2 Uch2-mCherry::ura4<sup>+</sup>* | 4B, S4D |
| 6938 | *h<sup>+</sup> Kan::P<sub>nmt41</sub>-slp1<sup>+</sup> SV40 P-GFP-atb2::LEU2* | 1D, 2D, S4D |
| 7091 | *h<sup>-</sup> Kan::P<sub>nmt41</sub>-slp1<sup>+</sup> SV40 P-GFP-atb2::LEU2*<br>*Uch2-mCherry::ura4<sup>+</sup>*<br>*Cut14-3XHA::NAT leu1-32* | 2D, S4C |
| 7026 | *h<sup>-</sup> Kan::P<sub>nmt41</sub>-slp1<sup>+</sup> cdc2(as)M17::Bsd*<br>*SV40 P-GFP-atb2::LEU2 Uch2-mCherry::ura4<sup>+</sup>*<br>*leu1-32 Cut14-3XHA::NAT* | 1A-1D, S1A-S1C |
| 7029 | *h<sup>+</sup> ppa2Δ::kanMX6 Kan::P<sub>nmt41</sub>-slp1<sup>+</sup>*<br>*SV40 P-GFP-atb2::LEU2 Uch2-mCherry::ura4<sup>+</sup>* | 2A, 2C, S3A, S3B |
| 7055 | *h<sup>+</sup> wee1Δ::kanMX6 Kan::P<sub>nmt41</sub>-slp1<sup>+</sup>*<br>*SV40 P-GFP-atb2::LEU2 Uch2-mCherry::ura4<sup>+</sup>* | 2A, 2B, S3A, S3B |
| 7083 | *h<sup>+</sup> ppa2Δ::kanMX6 Cut14-3XHA::NAT*<br>*Kan::P<sub>nmt41</sub>-slp1<sup>+</sup> SV40 P-GFP-atb2::LEU2*<br>*Uch2-mCherry::ura4<sup>+</sup>* | 2D |
| 6909 | *h<sup>+</sup> cdc2(as)M17::Bsd* | 2C, 4C |
| 7130 | h- *cdc2(as)M17:Bsd ppa2Δ::kanMX6 leu1-32* | 2C |
| 7125 | h- *cdc2(as)M17:Bsd cut14-IAA17-ura4+ ura4-D18* | 3C |
| 7127 | h+ *cdc2(as)M17:Bsd*<br>*ade6::ade6+-Padh15-skp1-OsTIR1-natMX6-Padh15-skp1-*<br>*AtTIR1-2NLS cut14-IAA17-ura4+ ura4-D18* | 3C |
| 7135 | h- *cdc2(as)M17:Bsd Cut3-3Pk::HPH*<br>*ura4+::cut3-3Pk,cut14-natMX lys3+::cnd3-BsdMX*<br>*his5+::cnd2-3XHA,cnd1-bleMX* | 4C |
| 6170 | h- *natMX-P<sub>nmt81</sub>-cut14-IAA-ura4+*<br>*SV40-GFP-atb2-LEU2 ade6::ade6+-Padh15-*<br>*skp1-OsTIR1-hph-Padh15-skp1-AtTIR1-2NLS*<br>*kan::Pnmt41-slp1+ ura4-D18* | 4A |
| 6819 | h- *Cut3-3Pk::HPH* | 4A |
| 6990 | h- *kan::P<sub>nmt41</sub>-slp1+ Cut3-3Pk::HPH*<br>*ura4+::cut3-3Pk,cut14-natMX*<br>*lys3+::cnd3-BsdMX his5+::cnd2-3XHA,cnd1-bleMX* | 4A |

**Table S2.** Plasmids used in this study.

| Plasmid | Description | Figure |
| --- | --- | --- |
| 2075 | Cut3-Pk3, Cut14 with *S. pombe* 5' and 3'UTRs in pUra4<sup>AfeI</sup>-natMX | 4A-C<br>S4C-D |
| 2206 | Cnd3 with *S. pombe* 5' and 3'UTRs in pLys3<sup>BstZ17I</sup>-BsdMX | 4A-C<br>S4C-D |
| 2207 | Cnd2-HA3, Cnd1 with *S. pombe* 5' and 3'UTRs in pHis5<sup>StuI</sup>-bleMX | 4A-C<br>S4C-D |

